# Survey of Australian STEMM Early Career Researchers: job insecurity and questionable research practices are major structural concerns

**DOI:** 10.1101/2020.02.19.955328

**Authors:** Katherine Christian, Carolyn Johnstone, Jo-ann Larkins, Wendy Wright, Michael R. Doran

**Author notes:** Corresponding Author: Katherine Christian or Michael R. Doran or. Author Contributions: K.C., C.J., J.L., W.W., and M.R.D. designed the research; K.C. collected the data; K.C., C.J., J.L., W.W., and M.R.D. analysed the data; K.C. and M.R.D wrote the paper; all authors edited and approved the submission.

## Abstract

We sought to understand the pressures on Early Career Researchers (ECR) in the Science, Technology, Engineering, Mathematics, & Medicine (STEMM) disciplines, collecting data from 658 ECRs working in Australia. Respondents indicated a “*love of science*”, but most also indicated an intention to leave their position. Decisions were primarily motivated by job insecurity (52%), while grievances included poor supervision (60%), bullying or harassment (34%), inequitable hiring practices (39%) and poor support for families (9.6%). A concerning rate of “*questionable research practices*” by colleagues (34.1% to 41.1%) was reported to have impacted ECR career advancement. Our study links recent reports that characterise the health of the research industry, providing direct insight from ECRs on job insecurity, workplace culture challenges, and the logical rise of questionable research practices. Internationally, nationally and institutionally the research community needs to improve job security (care for our people) and the quality of research data (our product).

## Introduction

Advances in Science, Technology, Engineering, Mathematics and Medicine (STEMM) have revolutionized virtually every facet of modern life, and further advances in the STEMM and/or STEM fields are assumed to underpin Australia’s future economic prosperity ^1^. Based on the merit of having a highly trained STEMM workforce, and on the service export value of education, Australia has expanded to become the largest provider of education to international students in the Organization for Economic Co-operation and Development (OECD) nations ^2^. These increases include postgraduate education, where the number of PhD completions in Australia has grown from 4,000 per annuum in 2000, to approximately 10,000 per year in 2019 (values include both domestic and international students, ^3^).

The proportion of PhD completions in Australia is slightly greater (1.17% of the working population) than the OECD average (0.99%), but is lower than the USA (1.78%) or Germany (1.38%) ^4^. Two international surveys conducted in 2015 ^5^ and 2017 ^6^ indicated that nearly 78% and 75% of PhD candidates, respectively, aspired to obtain a job in academia, despite the global lack of such job opportunities. The potential for disconnect between numbers of graduates and academic positions has been amplified by the doubling of PhD graduates numbers from 0.8% to 1.6% of the working population in the OECD between 1997 and 2014 ^7^. Not all PhD graduates need work in academia, however as the advanced industries that typically employ highly skilled workers are less developed in Australia than in many other countries such as USA or Germany ^8,9^, Australian graduates are more dependent on academia as an employer than in other markets ^3^. In Australia, the problem is acutely complex, with McCarthy and Wienk ^3^ noting in 2019 that there are not enough jobs in academia for all PhD graduates, and that the number of graduates has significantly outpaced academic jobs available in Australia since the mid-1990s.

A previous survey of Australian postdoctoral researchers identified that greater than half (52%) took their position hoping to transition to a full-time research role in academia ^10^. This survey captured feedback from a total of 284 postdoctoral researchers, of which approximately 80% were within 10 years of PhD completion (2–5 years post-PhD, 50%; 6–10 years post-PhD, 29%). The majority of respondents (54%) felt that structural, rather than personal limitations would prevent them from realizing a long-term research career. Respondents cited inadequate job security (37%), lack of funding (37%), lack of independent positions available (14%) and family or carer responsibilities (6%) as potential reasons for leaving academia. These previous data highlight that while Australian postdoctoral researchers work more than their contracted 38-hour week (on average), and desire to maintain a long-term research career, most appreciate that job access will limit this outcome.

In parallel to the international so-called “*glut*” of PhD graduates ^6,11^ and limited access to Australian research postdoctoral and academic positions ^10^, there has been the noted international rise of the reproducibility crisis ^12^. Estimates indicate that in 2018 there were 33,100 peer-reviewed English language journals, contributing approximately 3 million new articles per year to the literature, at an annual growth rate of 5-6% ^13^. The value of this output is contingent on the published data being accurate, and the published methods being reproducible. The reproducibility crisis (sometimes referred to as the replicability crisis) refers to the observation that many scientific studies are difficult or impossible to replicate or reproduce, including many studies reported in high impact journals. In a 2012 publication, Glenn Begley explained that before Amgen (Thousand Oaks, California) pursued a particular line of research, they would first attempt to reproduce the studies that underpinned the potential therapy ^14^. Of the fifty-three ‘landmark’ published studies they attempted to reproduce, the study’s findings were only validated in 6 (11%) cases. In 2016, Monya Baker detailed feedback from Nature’s survey of 1,576 researchers on the topic of data reproducibility ^12^, finding that pressure to publish and selective reporting, were perceived to contribute to greater than 60% of reproducibility problems.

Early career researchers (ECR) are a critical link between PhD programs and career researchers, and their well-being provides insight into the health of the industry. We surveyed STEMM ECRs in Australia to better understand the pressures impacting them and their career development. Data were collected from respondents employed in research institutions or universities via an on-line survey (n=658); surveys were developed based on previously published questions and through focus group discussions. We quantified satisfaction with work environments, likelihood of continuing to work in research in Australia, mentoring and career planning, and observation of questionable research practices. Our data provide a warning that the systemic pressures compromising Australian ECR training and career progression may also contribute to a decay in research quality. It is time to carefully consider if the STEMM ECR support and career advancement options are aligned with Australia’s scientific aspirations. As many of the documented pressures are common global problems, these data likely highlight important considerations relevant to the international research community.

## Results

Of the 658 respondents to our survey, 48% identified as being in the medical and health sciences. Recent data from the Australian Research Council (ARC) ^15^ indicates that 38.9% of Australia’s STEMM workforce is employed in the medical and health sciences (Table 1). Comparison of our survey demographics with this ARC data indicates that our sample and the target population were not statistically significant by discipline (chi square = 16.344, df = 9, p = 0.06), and our survey population can be considered representative.

**Table 1.** Percentage of Australian STEMM workforce, relative to percentage of survey responses segregated by discipline (n = 658). **Australian work force data sourced from ^15^.

| Discipline | **Percentage of Australian academic STEMM workforce | Percentage of respondents to this survey |
| --- | --- | --- |
| Mathematical Sciences | 3.8% | 2.8% |
| Physical Sciences | 4.3% | 8.1% |
| Chemical Sciences | 4.7% | 5.7% |
| Earth Sciences | 3.5% | 3.0% |
| Environmental Sciences | 3.2% | 4.0% |
| Biological Sciences | 12.6% | 20.9% |
| Agricultural and Veterinary Sciences | 4.5% | 1.4% |
| Information and Computing Sciences | 6.9% | 2.2% |
| Engineering | 15.4% | 3.6% |
| Technology | 2.1% | 0.8% |
| Medical and Health Sciences | 38.9% | 47.5% |

We attempted to identify workplace characteristics that influenced ECR job satisfaction and career progression. Our respondents almost universally noted their “*love*” of research and the job fulfillment it provides (Table 2). ECRs reported that they derived fulfilment from research, mentoring, teaching and the general sense that they are making a meaningful contribution to society.

**Table 2.** Quotes from Australian STEMM ECRs as to *Why they stay in science*.

| Quote number | Specific response |
| --- | --- |
| 1 | *I love figuring stuff out. I love inventing new ways to measure stuff.* |
| 2 | *I love it! I am passionate about my work and driven to make a difference. I will keep going as long as I can.* |
| 3 | *I love it. every day is different. I am inherently nosy so love that I can spend my day researching something which interests me.* |
| 4 | *I love my job - it doesn't feel like a job - I get to do what I enjoy. That said, the lack of job security and the challenges of having a family, buying a house and staying in the one city in Australia makes it difficult to imagine remaining in research/academia.* |
| 5 | *I love my job, and I want to make a difference.* |
| 6 | *I love my job, being able to develop new research questions and work with clinicians and patients. But I do not love the industry. The lack of job security, challenges in supporting a team, and constant pressure to do more as soon as you can is deeply problematic.* |
| 7 | *I love research and discovery, a core part of my identity is 'scientist'. I'm not sure who I would be outside academia.* |
| 8 | *I love research and I love teaching, and academia offers the opportunity for both of these. Improved job security would be the one key thing to improve my experience.* |
| 9 | *I love research and mentoring/training research students.* |
| 10 | *I love research and my research area, I want to help people through my science discoveries and the sharing of these results.* |
| 11 | *I love research and really believe that it can change clinical practice and therefore people's lives.* |
| 12 | *I love research and the people I meet and work with in the field.* |
| 13 | *I love research and understanding how new information fits into the bigger.* |
| 14 | *I love research, I love supervising students, I love writing papers, I love engaging the Public with Science.* |
| 15 | *I love research! No two days are the same and it is extremely rewarding. You have to celebrate the few good days you have (manuscript accepted, award at a conference, grant etc.). The opportunity to truly make a difference to the lives of people is what keeps me going!* |

We queried ECRs regarding satisfaction with their workplace culture. Academic workplace culture, which encompasses interactions between colleagues and professional norms ^16^, has evolved with corporate pursuits of universities and hypercompetitive funding environments ^17^. Figure 1a shows that 51.0% of respondents indicated that they were satisfied or very satisfied with their workplace culture, while a concerning 31.9% were somewhat or very dissatisfied with their workplace culture. Previous studies have identified diversity and inclusion as factors that have impact on senior academics’ dissatisfaction ^18,19^, including the career progression for female academics ^20-22^. However, in our survey of Australian ECRs, most identified as satisfied, or at least unconcerned, regarding discrimination with respect to age, gender, ethnic background or sexual orientation. Only 17.8% were dissatisfied with their workplace’s approach to diversity and inclusion (Survey question 31-13 (data not shown)). Our data, explored below, suggests that other challenges are dominant for this ECR cohort.

**Figure 1.**
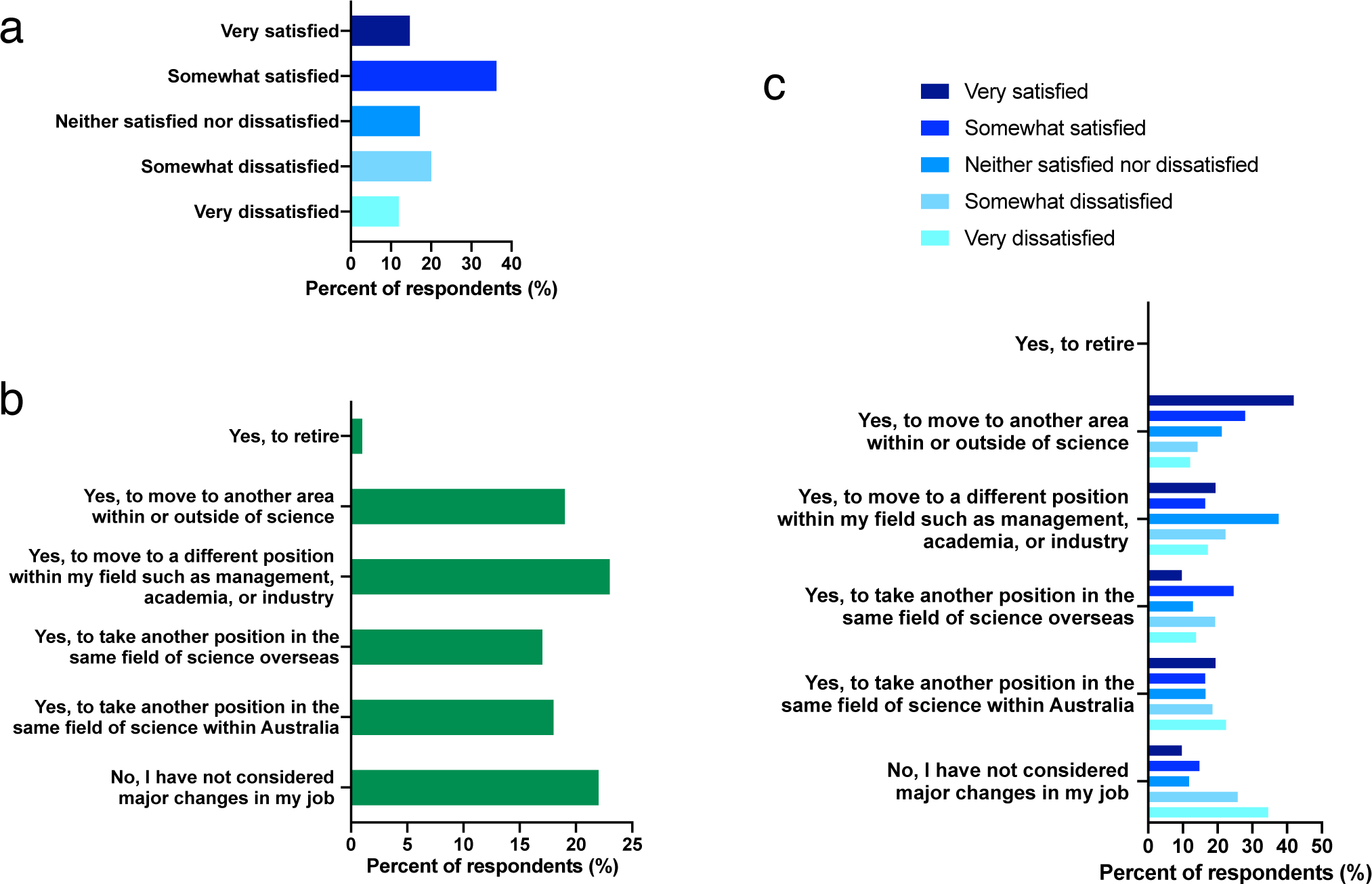
(**a**) Australian STEMM ECRs were asked to rate their overall satisfaction with their current work (Question 31-4 in survey, n = 566), (**b**) ECRs were asked if within the last five years they had considered any major career or position changes, and what these might be (Question 61 in survey, n = 471). (**c**) For those considering a major career or position change in the previous 5 years, we stratified responses from respondents based on satisfaction with their current position (n = 471).

We asked if ECRs had considered a major career or position change in the previous 5 years. The majority (78.3%) respondents had considered a major career change, while only 21.7% had not (Figure 1b). Many considered leaving academia all together (19.1%) or moving overseas (17.4%) in order to progress their career path. For each group of respondents that indicated that they had considered a major career change, we quantified how satisfied they were with their current work environment (Figure 1c). Interestingly, within the population of ECRs who had not considered a career change, the largest group (34.5%) were dissatisfied with their current workplace. By contrast, within the population of ECRs who indicated that they had recently considered moving to another area within or outside of science, the largest group (41.9%) were very satisfied with their current workplace. These data might suggest that there are populations of ECRs who are unhappy in their current workplace, but feel trapped, while there is another population of ECRs who are very happy in their current workplace, but feel changing jobs would be beneficial. More generally, ECR’s satisfaction with their current position does not appear to correlate with their consideration of major career changes or moves.

Workplace and career progression challenges are described in Table 3. Data were sorted based on gender in Table 3a, and subsequently sorted based on appointment type in Table 3b. Appointment types were categorized as “*research only*”, “*research and teaching*”, or “*clinician researcher*”. Those with a teaching or clinical appointment are likely to be less dependent on research funds for their salary, and thus their perspectives may differ. Greater than 50% of both male (52.4%) and female (63.8%) ECRs indicated that they felt they had been negatively impacted by a lack of support from institutional leaders (Table 3a). Female ECRs indicated higher rates of inequitable hiring practices (40.0% females versus 35.4% males) and harassment from those in a position of power (31.7% females versus 25.9% males). Interviews and a focus group conducted in addition to this survey, as well as survey responses, suggest instances where senior academics (both male and female) were regarded as bullies. When asked if they feel safe in the work environment, overall 12.5% felt unsafe with an unexpected bias of males (15.6%) to females (11.0%) reporting this problem (Table 3a).

**Table 3.**
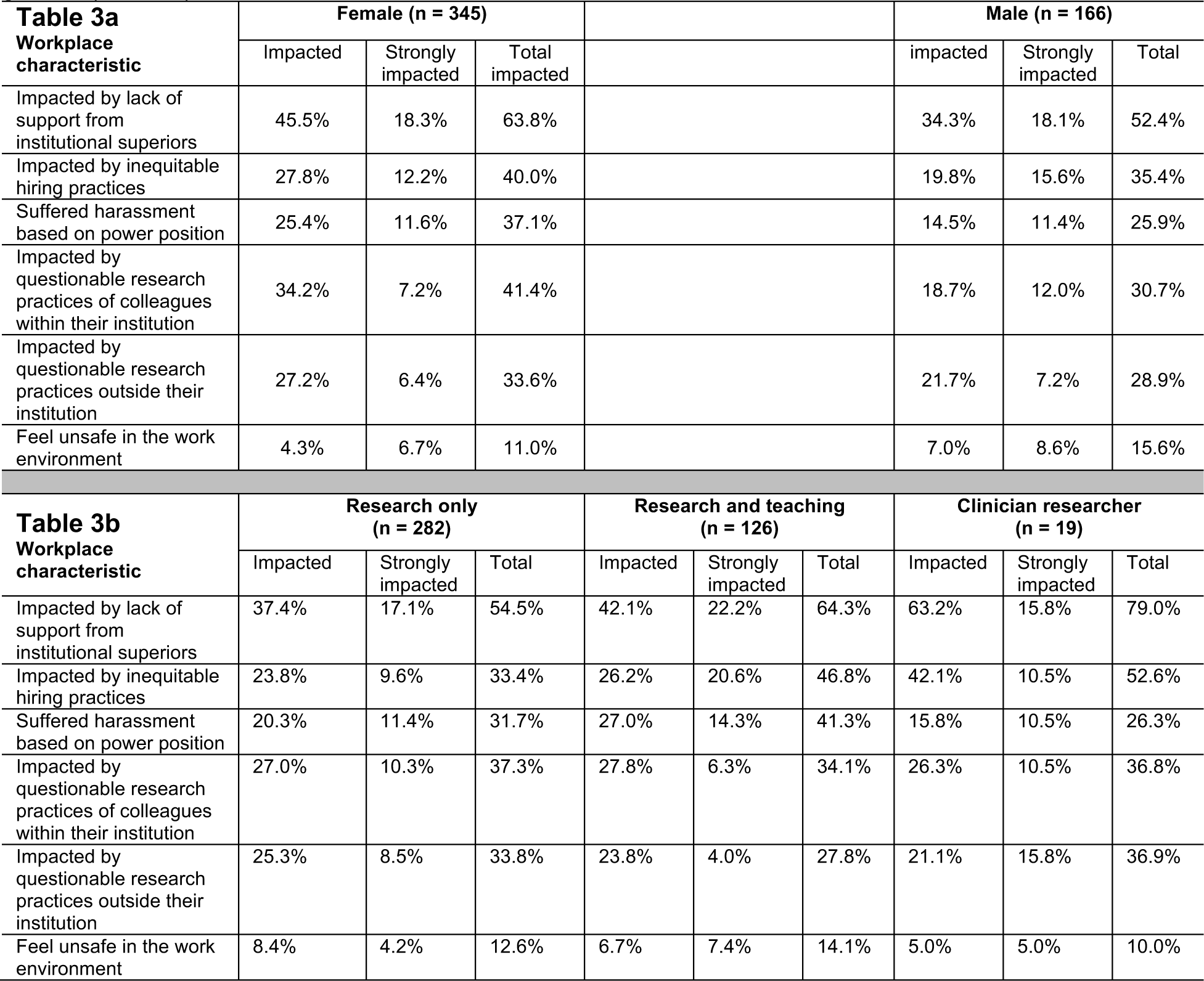
(**a**) Factors that impacted on ECR job satisfaction and/or career progression, analysed with respect to ECR appointment type (n = 517). (Teaching only and “Other” responses are omitted from this Table 3a). (**b**) Factors that impacted on ECR job satisfaction and/or career progression, analysed with respect to gender (n = 511).

| **Table 3a**<br>Workplace<br>characteristic | Female (n = 345) |  |  | Male (n = 166) |  |  |
| --- | --- | --- | --- | --- | --- | --- |
|  | Impacted | Strongly impacted | Total impacted | impacted | Strongly impacted | Total |
| Impacted by lack of support from institutional superiors | 45.5% | 18.3% | 63.8% | 34.3% | 18.1% | 52.4% |
| Impacted by inequitable hiring practices | 27.8% | 12.2% | 40.0% | 19.8% | 15.6% | 35.4% |
| Suffered harassment based on power position | 25.4% | 11.6% | 37.1% | 14.5% | 11.4% | 25.9% |
| Impacted by questionable research practices of colleagues within their institution | 34.2% | 7.2% | 41.4% | 18.7% | 12.0% | 30.7% |
| Impacted by questionable research practices outside their institution | 27.2% | 6.4% | 33.6% | 21.7% | 7.2% | 28.9% |
| Feel unsafe in the work environment | 4.3% | 6.7% | 11.0% | 7.0% | 8.6% | 15.6% |

| **Table 3b**<br>Workplace<br>characteristic | Research only (n = 282) |  |  | Research and teaching (n = 126) |  |  | Clinician researcher (n = 19) |  |  |
| --- | --- | --- | --- | --- | --- | --- | --- | --- | --- |
|  | Impacted | Strongly impacted | Total | Impacted | Strongly impacted | Total | Impacted | Strongly impacted | Total |
| Impacted by lack of support from institutional superiors | 37.4% | 17.1% | 54.5% | 42.1% | 22.2% | 64.3% | 63.2% | 15.8% | 79.0% |
| Impacted by inequitable hiring practices | 23.8% | 9.6% | 33.4% | 26.2% | 20.6% | 46.8% | 42.1% | 10.5% | 52.6% |
| Suffered harassment based on power position | 20.3% | 11.4% | 31.7% | 27.0% | 14.3% | 41.3% | 15.8% | 10.5% | 26.3% |
| Impacted by questionable research practices of colleagues within their institution | 27.0% | 10.3% | 37.3% | 27.8% | 6.3% | 34.1% | 26.3% | 10.5% | 36.8% |
| Impacted by questionable research practices outside their institution | 25.3% | 8.5% | 33.8% | 23.8% | 4.0% | 27.8% | 21.1% | 15.8% | 36.9% |
| Feel unsafe in the work environment | 8.4% | 4.2% | 12.6% | 6.7% | 7.4% | 14.1% | 5.0% | 5.0% | 10.0% |

Particularly concerning was that the number of female and male ECRs who identified that they themselves, or their career advancement, had been impacted by *questionable research practices* within their institution (41.4% of females and 30.7% of males) or external to their institution (33.6% of females and 28.9% of males). While some respondents would have been cautious not to reveal specifics regarding questionable research practices, even in a confidential survey, a number of comments did provide reasonably detailed examples of concerning behaviour (Table 4).

**Table 4.** Quotes from Australian STEMM ECRs regarding *questionable research practices* (from surveys and interviews).

| Quote number | Specific response |
| --- | --- |
| 1 | *....the bullying and stuff came to a head and the scientific work was looked at because this person had brought up kind of bullying and harassment allegations against the supervisor. So they in turn looked at the work that this person had been doing and they'd been falsifying...* |
| 2 | *Lack of funding and the need to 'sell' your research, often leads to many researchers fabricating and embellishing data. This leads to the inability of genuine researchers to replicate findings, wasting precious time and resources, giving up and then their contracts not being renewed because the boss doesn't get the 10 publications per year they demand.* |
| 3 | *I believe that the whole Academia environment is corrupted and has lost its true vision. The lack of funding is making researchers to sometimes make-up data to get grants or to publish meaningless papers just for the sake of raising the numbers.* |
| 4 | *being used by post docs and high level senior researchers' who take credit for your research work ideas and use info in your recruitment applications unethically for themselves...bias recruitment towards international students and overseas post docs who are extremely competitive and who want to get permanent residency and who also bully harass local students and researchers' to take over their research and jobs.* |
| 5 | *...what they wanted to see result-wise wasn't what I was seeing. And so I was being accused of misconduct because I wasn't seeing what they wanted me to see, and I wouldn't change that.* |
| 6 | *Not saying, 'do this' but pressure to – if something were to fail to almost keep saying, 'Do it again, do it again, do it again, do it again' in order to get you to make it work. And those people have just said, 'No, it doesn't and I'll spend the whole year repeating it but it's not going to change the outcome'.* |
| 7 | *Q But are they getting their names on because they've actually been involved? Are we flouting the convention here?*<br>*A They haven't done anything.*<br>*Q So his investment in them is...*<br>*A Is purely so they can get grant funding through having papers.* |

When the data from Table 3a was re-sorted based on appointment type, it was possible to estimate the influence that different appointments and contract stability may have on ECR job satisfaction and/or career progression (Table 3b). We only had a small number of clinician researchers (n = 19). Such researchers, in most cases, rely primarily on their clinical appointment as a source of income, and so are potentially less sensitive to job insecurities felt by research only ECRs. The majority of clinician researchers (79.0%) reported having been impacted by lack of support from institutional superiors. Only two (10.5%) of clinician researcher ECRs reported feeling unsafe at work, suggesting that clinical job security may protect from such circumstances. The frequency that *questionable research practices* had negatively impacted their careers declined incrementally from those who were research-only (37.3% internally and 33.8% externally), clinician researchers (36.8% internally and 36.9% externally) and research and teaching (34.1% internally and 27.8% externally). These data suggest that greater research time commitment may increase the frequency of exposure to *questionable research practices*, but that the stability associated with salary funding from a teaching or clinical position does not obscure the perception that this is a major problem.

When data were delineated based on years post PhD, those who were greater than 4 years post PhD were less satisfied than those who were 4 years or less post PhD (55.7% vs 66.9%). Similarly, those who were greater than 4 years post PhD tended to indicate a higher frequency of being negatively impacted by lack of support from institutional supervisors (67.5% vs 54.1%), questionable research practices of colleagues within their institution (41.3% vs 35.8%), and harassment based on power position (36.7% vs 31.1%). These more senior postdoctoral researchers more often indicated that if they had their time again they would not become an academic (38.1% vs 28.4%), and were more likely to not recommend science as a career to a young person (78.2% vs 53.5%).

We observed that the academic culture promotes a perceived *need* to relocate during the ECR years, and that many ECRs who wished to remain in academia considered moving as part of their career development process. To better understand this phenomenon, we asked more detailed questions regarding decisions to move. Our data indicates that moves to new institutions can be stressful (Table 5). Relocation, often to new countries, is frequently made without financial compensation for the relocation and can be challenging for families, and for careers. A 2020 editorial in Science describes the struggles of a tenure-track academic living in the USA on a work visa, and his inability to gain bank finance approval to purchase a home ^23^. While a tenure-track academic can make long-term decisions, this is virtually impossible for many ECRs. Most (68.1%) respondents reported that they had already changed location in order to advance their careers. Of these, 28.6% of ECRs had moved once, 20.1% had moved twice and 19.5% had moved more than twice. Commonly expressed consequences, noted in interviews and in text-based responses were that relocation was associated with stress, separations from family, loss of support network, personal cost and loss of career momentum.

**Table 5.**
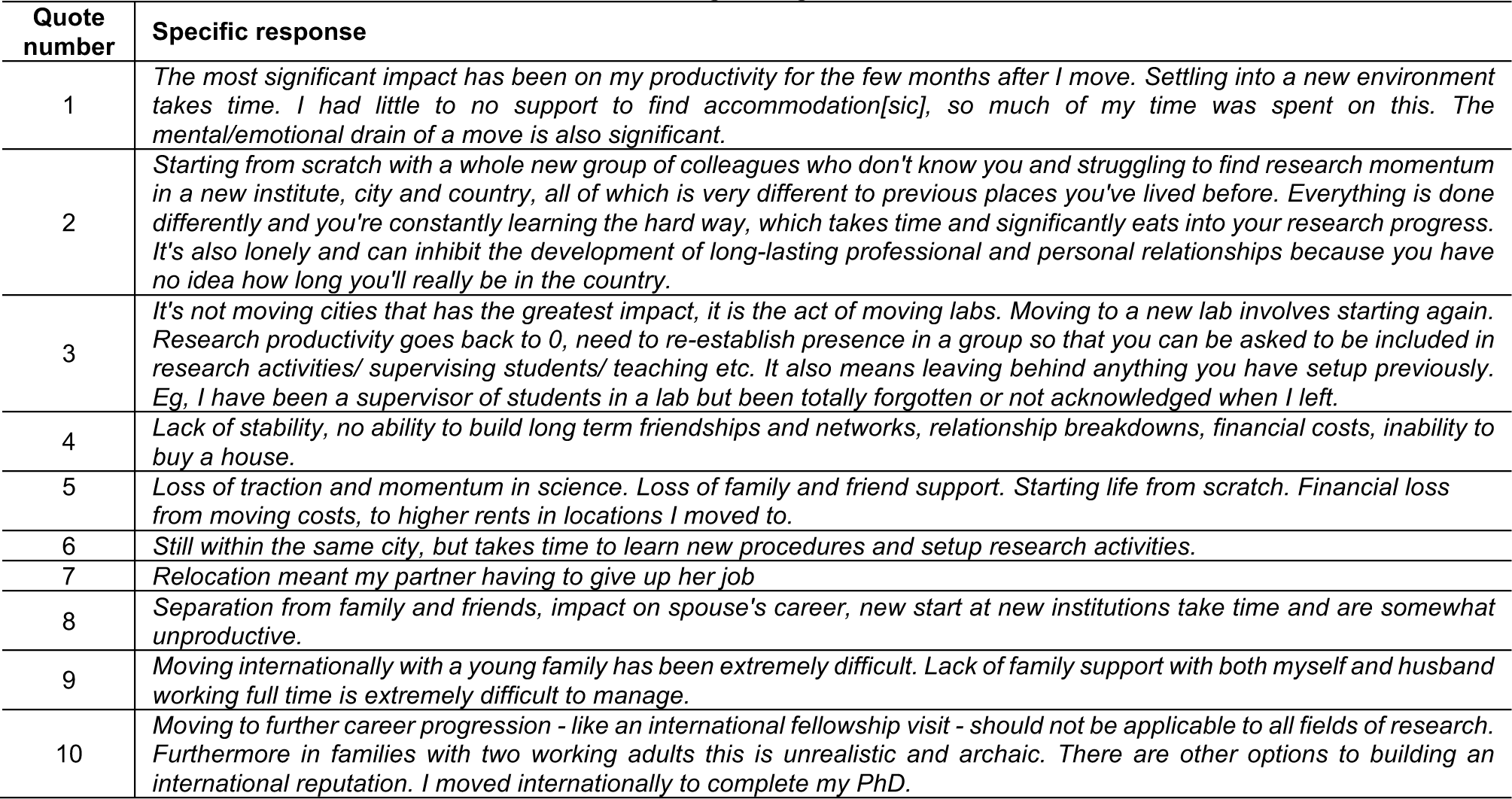
Quotes from Australian STEMM ECRs regarding the stress of relocation.

To better understand ECRs concerns regarding support from institutional leaders, respondents were asked to describe their mentorship and career guidance. A definition of a of mentor was provided with the questions: “A mentor is someone who is there to assist you achieve your personal, academic and career exploration goals. This person is not necessarily your supervisor”. In our survey, 61.9% of ECRs reported having a mentor, while 38.1% did not. We asked ECRs to indicate what aspects of mentoring they valued most, and these data are summarized in Table 6A. ECRs valued advice on career decisions (81.7%) as the most important contribution from mentors. This was followed by integration into networks (77.2%), and direct influence on their gaining employment (56.7%). Ranked less significant, but still important, were skill training on methodologies (60.3%), fundraising (50.8%), and scientific writing (59.7%). Of those with a mentor, the quality of the mentoring was often described as inadequate, and some indicated that they paid for external mentoring. From the survey data (n = 322), those who did receive mentoring (Question 44 of our survey) described it as follows; 15.1% neutral, 7.5% not beneficial, 32.8% highly beneficial, or 44.6% beneficial.

**Table 6.** (**a**) We asked ECRs to indicate how much value they placed on different aspects of mentoring or supervision (n = 481 respondents). (**b**) We asked ECRs that participated in performance reviews to indicate which aspects of the review process they valued (n = 322 respondents)

| **Table 6a Value of mentoring or supervision contribution** | **Unimportant** | **Neither important nor unimportant** | **Important** |
| --- | --- | --- | --- |
| Advice on career decisions | 6.4% | 18.3% | 81.7% |
| Introduction to important networks | 6.8% | 22.8% | 77.2% |
| Attain a position via direct intervention | 17.0% | 26.3% | 56.7% |
| Skill training: methodology | 17.0% | 22.6% | 60.3% |
| Skill training: fundraising | 21.4% | 27.8% | 50.8% |
| Skill training: scientific writing | 17.8% | 22.4% | 59.7% |

| **Table 6b Rate benefit of mentoring activity** | **Not at all useful, or not useful** | **Neither useful, nor not useful** | **Useful, or very useful** | **NA** |
| --- | --- | --- | --- | --- |
| Overall | 41.6% | 13.0% | 43.8% | 1.6% |
| In reviewing your personal progress | 28.0% | 13.4% | 57.1% | 1.6% |
| In identifying your strengths and achievements | 29.9% | 17.8% | 50.7% | 1.6% |
| Helping focus on career aspirations and how met by current role | 33.3% | 14.6% | 50.4% | 1.6% |
| For you to highlight issues | 35.1% | 18.5% | 44.2% | 2.2% |
| In leading to training or professional development opportunities | 48.0% | 18.8% | 31.6% | 1.6% |
| In leading to changes in work practices | 58.2% | 22.3% | 16.6% | 2.8% |

With respect to supervision, as opposed to mentoring, only 68.3% of respondents had a performance review in the past two years, indicating that 31.7% had not. While half of the 31.7% respondents with no performance review indicated that they had recently been appointed or were on probation (not unusual in an environment where short term contracts are commonplace), the other 50% had not been offered a review. Many who did have a performance review did not find the process useful (41.6%, Table 6B). There was no opportunity given to provide an explanation for these answers, however respondents identified the primary utility of performance reviews as being (1) a review of personal progress (57.1%), (2) identifying strengths and achievements (50.7%), (3) help focusing on career aspirations (50.4%), and (4) to highlight issues (44.2%). ECRs identified performance reviews as least useful in leading to changes in their work practices. Given that performance reviews are often used to influence work practices, it is useful to know that this process is frequently viewed as ineffective.

Finally, we circled back and considered if the positions ECRs held were similar to what they had anticipated, and if they intended to remain in or leave these positions (Figure 2a). Relatively few (14.5%) found their current position to better or much better than expected. Additional direct comments from respondents to this survey question can be found in Supplementary Table 1. Regardless of their perception of the position, many ECRs indicated their intention to leave. There was a trend (regression analysis, p = 0.0234) indicating a greater bias to leave the position depending on how it had met expectations (Figure 2b). However, even in instances where the current position was much better than expected, nearly 40% more (61.5%) ECRs intended to leave the position rather than remain (38.5%). As most ECR positions are short-term contracts, including those supported by “soft money” (where all expenses for that researcher, including salary, are covered by fixed-term grants), it might be rational to expect to have to leave a position even if the position had met or exceeded expectations. If ECRs were to leave their current academic position, we asked what the primary motivation would be (Figure 2c). Cumulatively, two of the possible responses, lack of funding (28.2%) and job insecurity (48.9%), accounted for 77% of likely motivations for ECRs leaving their current position. Establishing an independent research group is the goal of many ECRs. Lack of independent positions was cited as the motivation 11.8% of ECRs would use to justify leaving their current position. While in Table 3 many respondents list poor institutional support as problematic, only 1.4% of respondents cite interpersonal relationships with their supervisor as a potential motivation for leaving their current position. We found that Family/Carer responsibilities was cited by 9.6% of ECRs as a reason to exit academia. Similar to a previous Australian Postdoctoral Researcher survey ^10^, the burden of Family/Carer responsibilities is heavy on both male and female ECRs, suggesting that young parents (male or female) and their families are not sufficiently accommodated by the current system. In interviews, we did identify young mothers on parental leave struggled to continue to run their laboratories, knowing that their staff depend on them, and continued to write publications while on leave out of fear of falling behind. Quotations in Table 7 provide insights into stresses felt by Australian STEMM ECRs; we leave the quotes to speak for themselves.

**Table 7.** Quotes from Australian STEMM ECRs regarding stresses in the current system (explanations offered for responses to Question 73).

| Quote number | Specific response |
| --- | --- |
| 1 | *I just find the other aspects of the job and the pressure to perform very difficult. I feel like there is a big clock ticking, and my productivity is always being judged relative to the steady ticking of that clock regardless of the ups and downs and other life circumstances.* |
| 2 | *I just wish that the environment didn't feel so pressured and competitive. I have seen so many great ECRs leave research because of the challenges of finding work, meeting expectation, attracting grants. I think the field is too competitive and does not take care of our ECRs and we are poorer for it.* |
| 3 | *I am currently looking outside academia to get away from the culture of harassment... it takes too much of a toll on my health... but I would stay in academia if I were to find a position that didn't subject me to harassment by a supervisor.* |
| 4 | *The pressure of continual high performance and attracting funding would likely be too much and I'd opt for something easier and less stressful.* |
| 5 | *Job security is based on churning out a large quantity of publications, regardless of quality. Three-year fixed-term contracts are very short. In the first 2 years, I focus on my research, however, in my final year, I am thinking about where I am going next. It takes a lot of time and effort to find something else within the research field. I find having an "exit strategy" important.* |
| 6 | *Having said that, the pressures of the job have considerably increased in the last ten years and the general expectation is that you should work outside normal working hours, without getting paid extra... And that being able to work in academia is a privilege, so one should do whatever it takes to continue in Academia. In my opinion this is a very distorted and dangerous vision, which puts lots of pressure on ECRs, in particular women who are usually starting families at this stage in their careers.* |
| 7 | *At the point of my career, where I am trying to expand my group to potentially have an independent research group, the stresses around funding are a considerable issue for me (as for everyone else, probably). While I have been relatively successful with funding, I feel the pressure of having to support not only my own research, but also the research of those who work with me, and that holds me back from pursuing opportunities that are available to me as I don't want my group to expand too quickly. It also means that I put up with being paid on a lower pay scale than I should be, rather than going for promotion, because I want to conserve funding. This is certainly a constraint on my ability to expand my career prospects.* |
| 8 | *The personal toll it takes to have an academic position is immense. The job insecurity, being unable to plan for anything beyond 1-maybe 2 years is debilitating. Constantly responding to this opportunity, and that opportunity, doing good clever work and being available at all times is tough beyond measure. Not knowing if all this personal sacrifice and tough hard work are even going to be worth it is downright demoralizing. It might all work out, and it might not - but when do you pull the pin??* |
| 9 | *This way I hardly see a way to progress without affecting my mental, physical and emotional health due to the day to day stress.* |
| 10 | *Mental health of ECRs is overlooked and the universities treat us as second class employees that are disposable.* |

**Figure 2.**
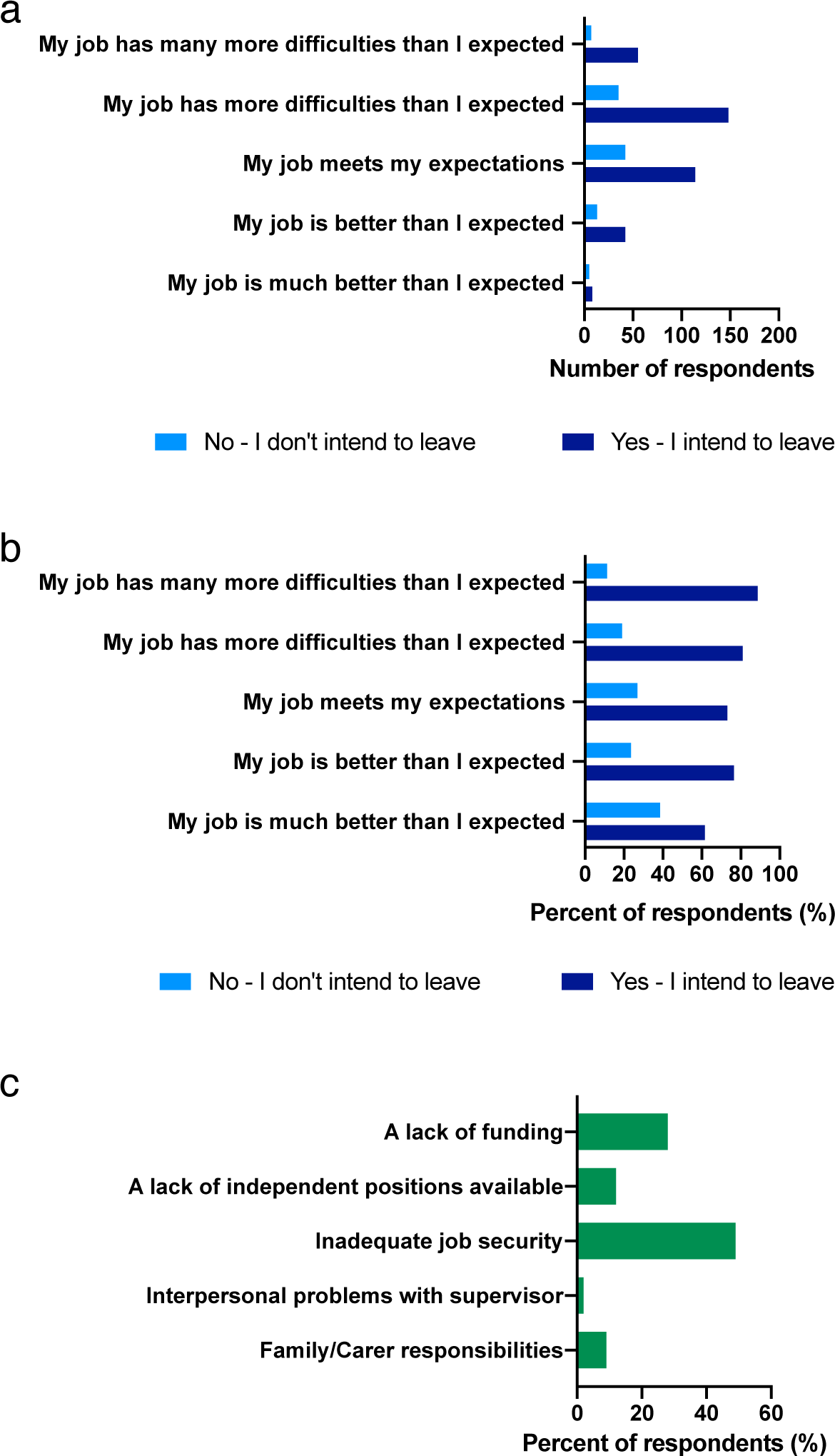
ECRs expectations of their current position, and their intention to leave. (**a, b**) Survey Question 73; *How does your job as an early-career researcher meet your original expectations?* (n = 469), and ECRs’ intention to leave or remain in that position. (**a**) Data shown as raw number of respondents. (**b**) Data shown as percentage of each group of respondents. Note correlation between job expectation and intention to leave (n = 469, regression analysis, p = 0.0234). (**c**) These data outline likely reasons for why ECRs would consider leaving a career in research (Question 67 in survey, n = 425, note that 38 answered other and are not accounted for in this graph).

Given the many challenges faced by ECRs, their persistence in their endeavours to remain in the academic research workforce is impressive. However, their perceived commitment to academia in Australia may be confounded by limited number of alternative (perceived and actual) employment opportunities outside of academia. A number of comments made by ECRs (Table 8), indicate that they consider themselves to be inadequately trained for alternative careers, that there are limited alternatives available, or that they regard leaving academia as a failure.

**Table 8.** Quotes from Australian STEMM ECRs regarding why they do not exit from academia, and their fears regarding employment outside of the academic workplace.

| Quote number | Specific response |
| --- | --- |
| 1 | *Because it took me so long to earn my PhD, not using it now would seem like a waste. Also, I don't know what else I am qualified to do.* |
| 2 | *I didn't know what the other options were or how to pursue them.* |
| 3 | *I enjoy science. I feel like leaving would be a failure. I try to continue / stay alive until that failure happens.* |
| 4 | *I've spent 10 years training to be an academic. I want to be an academic, but it seems it just isn't my choice at the end of the day. I'll stay until I am no longer competitive. I am keeping my eyes open and looking at other opportunities but so far no one wants me outside academia either.* |
| 5 | *I have no experience in any other field and I'm already 35 years old.* |
| 6 | *I have no skills in anything else.* |
| 7 | *After 13 years at university, a divorce, my body and mind falling apart, and pulling myself up from grinding childhood poverty and abuse there isn't anything else I feel that I am qualified to do. I am really good at my job yet overqualified and not healthy enough to do anything else. I am stuck here.* |
| 8 | *I also cannot imagine working in another environment, I actually don't know what other options are available and whether these would be fulfilling.* |
| 9 | *After spending all that time studying, it seems like a waste to move into a different field.* |
| 10 | *Fear, I have bills to pay and not a lot of financial support. If I leave I'm afraid I won't get another job* |
| 11 | *I constantly think about leaving academia/research (from necessity not choice) but don't know how and am not qualified for any other jobs.* |
| 12 | *I applied for a range of academic, research, technical, managerial and information technology positions that had more clear future prospects. The private sector did not seem interested in my academic background. Retraining is an option, but is likely to come at a significant cost to my current academic path.* |

## Discussion

A requisite attribute for a career academic is that individuals successfully navigate the process of being an ECR. The ECR career stage must necessarily function to identify and promote talent, as well as provide the professional training required to lead future research projects. We queried Australian ECRs from the STEMM disciplines with the goal of understanding their motivations, their challenges, and areas where their training environment could be improved.

Previous studies identified that academics loved their work and realised intellectual satisfaction, but were frequently discontented with their own institution and wonder if they would be happier somewhere else ^24,25^. Australian ECRs in our survey overwhelmingly and repeatedly indicated that they “*loved*” their work. However, as reported in Results, only 51.0% of ECRs indicated that they were satisfied with their workplace culture, and a concerning 31.9% indicated that they were somewhat or very dissatisfied. More than half of ECRs (male, 52.4%) and (female, 63.8%) felt they had been negatively impacted by a lack of support from institutional leaders. Female ECRs indicated experiencing higher rates of inequitable hiring practices (40.0% females versus 35.4% males) and harassment from those in a position of power (31.7% females versus 25.9% males) than their male counterparts. In contrast, more males (15.6% males versus 11.0% females) felt unsafe in their work environment.

Many ECRs indicated that they did not have a mentor (38.1%) or performance review (31.7%). Superficially, these data suggest that allocation of a mentor and performance review would lead to considerable improvements. However, a number of respondents (41.6%) indicated that they did not find the performance review useful. When mentoring and reviews were provided, ECRs valued career advice most (81.7%), followed introduction to important networks (77.2%), and the capacity of their mentor to directly help them find employment (66.7%). Ranked less significant, but still important, were skill training on methodologies (60.3%), fundraising (50.8%), and scientific writing (59.7%). These data may seem surprising, but a previous surveys of Australian postgraduate researchers found that the quality of supervision did not positively influence initial job attainment, but that “*nurturing networking and careers advice*” did ^26^. This pattern may remain robust in the Australian STEMM ECR cohort, where “*who you know”* could play a significant role in employment outcomes. Our data suggest that ECRs believe this is a factor, many report being impacted by inequitable hiring practices (40.0%, females and 35.4%, males). Job stress in the sector is likely causing similar patterns to evolve in jurisdictions around the world (see discussion on social networks and so call “gate keepers” and academic recruitment ^27^).

We do not dismiss the value of good mentoring and recommend that group leaders consider investing time into training and mentoring strategies (suggested reading: *Nature’s guide for mentors* ^28^). A recent paper in Nature Communications highlights how ECRs who co-author publications with highly-cited scientists have greater probability of repeatedly co-authoring additional publications with top-cited scientists, and, ultimately, a higher probability of becoming top-cited scientist themselves ^29^. While this does not directly constitute mentorship, it does provide an indication of the value of being able to follow or mimic an established research leader. The majority of ECR respondents identified mentorship on methodologies (60.3%), fundraising (50.8%), and scientific writing (59.7%) as important.

The value of any research is underpinned by the assumption that it is performed to a high standard and reported honestly. We consider the most concerning of all of our results to be the high rate at which ECRs (41.4% of females and 30.7% of males) claimed that questionable research practices within their institutions had negatively impacted their careers. Note that we did not define “questionable research practices” in our survey. In addition to fraud, in 2012 John *et al*., popularised the notion that *questionable research practices* included less egregious practices, such as selectively reporting studies that “worked”, rounding down p-values or failing to report all conditions ^30^. They concluded that the rate of these less egregious questionable practices was high, and that the high rates of these behaviours could be more damaging to the academic enterprise than more serious cases of fraud, which occur at lower rates^30^. Given the very high stress on individual ECRs and on the system, it is rational to expect that rates of questionable research practice could be on the rise. In 2005, Dr. John Ioannidis reasoned that “*most published research findings are false”*, discussing the influence of data selection bias and financial pressures on data interpretation and reported outcomes ^31^. Global research pressures have not declined since 2005, and in 2016 Nature published the results from a survey of 1,576 researchers on the topic of data reproducibility and the so-called reproducibility crisis ^12^. The Nature survey found that pressure to publish and selective reporting were perceived to contribute to greater than 60% of reproducibility problems. Our interpretation of survey data we collected, where ∼35% of respondents indicated that questionable research practices had impacted their careers, is that the full extent of known misconduct or data reproducibility problems is likely underestimated. Given that ECRs are both sufficiently trained to identify problems, and often in the laboratory enough to observe these problems, concern from this cohort should be viewed as genuine.

We highlight the need for institutional and national consideration regarding how pressures are playing out in the Australian STEMM research eco-system. We do not blame institutions or individual ECR mentors for these problems. Few ECRs (1.4%) indicated that they would leave their current position because of poor interpersonal relationships with their supervisor. Rather, we consider that the challenges experienced by Australian STEMM ECRs reflect systemic problems. Most ECRs (78.3%) had considered a major career change in the past five years, including leaving academia all together (19.1%) or moving overseas (17.4%). If ECRs left their current positions it would be primarily because of lack of funding (28.2%) and job security (48.9%). When the ECR responses were delineated based on years post PhD, those who were greater than 4 years post PhD were less satisfied than those who were 4 years or less post PhD. Our observations parallel a previous study that observed that job satisfaction was greater for those who had more recently started their first postdoctoral appointment ^32^. Within our cohort, senior ECRs more often indicated that this was not a good time to be in science, and were less often willing to recommend science as a career.

Both male and female ECRs are concerned about parental/carer responsibilities, knowing that delayed research productivity could compromise their career prospects. Men are more concerned about this than women, possibly reflecting recent efforts to accommodate mothers, but not necessarily families. It is common vernacular to say that “*ECRs are the future”*. If this is factually true, then are we content with how we are shaping this future? We suggest that this survey data provides reason to be concerned. Australia needs to invest in this future, but not at the expense of individuals. There is a disconnect between the current STEMM degree and postgraduate degree completion rates and workforce need for these graduates ^3^. Not all PhD graduates need necessarily work in academia, but employability outside academia will influence the dependence of PhD graduates and ECRs on academia as an employer. The weak development of advanced industries in Australia, relative to other parts of the developed world ^8,9^, has broad ramifications for the STEMM workforce which has expanded through the recent Australian education boom ^2,3^. In 2018 Australia spent 1.88 per cent of GDP on research and development, well below the OECD average of 2.38 per cent ^33^, whilst training a record numbers of PhDs ^3^.

Challenges for researchers are not isolated to Australia. In January 2020, the Wellcome Trust reported on results of a survey of 4,267 researchers, mostly from the UK ^34^, and a summary was published in Nature ^35^. Their data parallels many of our observations. While 84% of researchers were proud to work in the research community, only 29% felt secure in pursuing a research career. In the Wellcome Trust survey, 23% of junior researchers and students suggested that they had felt pressured by their supervisor to produce a particular result ^34^. Across the whole respondent population, 43% believed that their workplace puts more value on meeting research metrics, rather than on the quality of the research. Our survey went a step further, with ∼35% ECRs suggesting specifically that questionable research practices by colleagues had directly impacted their careers. It is clear that these are global challenges, that will require intervention at all levels of the research community.

Managing research training, careers and scientific output will require structural changes at the international, national and institutional level. Internationally, the scientific community needs to: (1) Think more critically about experimental design, data interpretation and statistical analysis (see statistical commentary here ^36^), and (2) have publications outline the limitations of their studies, including both scientific and practical translational limitations. These two changes could counter tendencies to overinterpret data or to hype outcomes. Nationally, Australia should consider: (1) An increase in GDP expenditure on research and development to align with the OECD, (2) trim PhD completion numbers to better align with current workforce demands, (3) distribute limited research funds through smaller packets that nurture a larger number of ECRs, recognizing that innovation and innovators are rare, and that time is required to test ideas and develop gifted researchers, and (4) establish an independent research ombudsman to oversee research integrity issues (need for an independent research ombudsman has been discussed previously (^37^ and ^38^). At the institutional level, around the world, the STEMM ECR research environment could be improved by: (1) training mentors to manage ECR career development, (2) aiming to provide greater career stability, (3) developing skills training programs that prepare PhD candidates and ECRs for employment outside of academia for when long-term academic employment is not viable, and (4) supporting the development of a research culture that counters questionable research practices by encouraging all academics to ask questions, challenge hype, and report honestly. As a community we need to work to improve job insecurity (take care of our people) and the quality of research data (our product).

## Materials and Methods

This research project explored challenges faced by early-career researchers (ECRs) in the sciences at universities and at independent research institutes in Australia. The primary research questions from which the survey questions were derived were; (1) What are the relationships between ECR job satisfaction or dissatisfaction and their likelihood of staying in science? (2) What are the principal factors that shape the ECR experience of various cohorts in the sciences in Australia? (3) What are the motivations for ECRs leaving the sciences? and (4) What are the specific features of the experiences and environment of those ECRs who remain in the sciences? The definition of “early career researcher” for the purpose of this project included holding a PhD or equivalent, awarded no more than ten years prior and employment in an Australian university or independent research institute in a STEMM discipline.

### Ethical Approval

This study has been conducted according to the guidelines of the ethical review process of Federation University Australia and the National Statement on Ethical Conduct in Human Research (Approval Number 18/139A) ^39^.

### Survey

Survey questions are included in the Supplementary Data Section (Supplementary Table 2). Quantitative data was collected from 658 respondents in an on-line survey of ECRs working in a scientific environment in universities and research institutes across Australia. The questionnaire for the survey was developed by first compiling questions, often used in a broader or international context, from research literature including questions from Australian Council of Education Research, The EMCR Forum (part of Australian Academy of Science) ^40^, Federation of Australian Scientific and Technological Societies (FASTS), Global Young Academy, National Science Foundation, Nature and Vitae ^10,41,42^ in order to cover all the themes identified in the literature as matters relating to job satisfaction or dissatisfaction. Some additional questions were created if no suitable question was identified elsewhere. Questions were combined and modified to create a question bank for this survey relevant to the research questions and the Australian context and further informed by data collected from a focus group of ECRs, after which the survey was pilot tested. Matters investigated include inequity, bias or discrimination with respect to age, gender, sexuality or race, inequitable hiring practices and harassment based on different power positions, mentoring and supervision, career planning, training and professional development and work life balance. The data from these questions were supplemented by questions seeking demographic information which included the institution type, research discipline, country of origin, family situation and work arrangements.

The invitation to take part in the survey was distributed via email after direct contact with the institutions, via social media or “umbrella groups” such as EMCR Forum ^40^ and The Australian Society for Medical Research (ASMR, ^43^) with members or affiliates drawn from the STEMM community who were likely to include the target group.

A focus group discussion attended by seven ECRs on January 30, 2019 evaluated the questionnaire prior to the survey and participants in the focus group offered additional insights. A pilot study (n=22) permitted testing for understanding and clarity and to check for technical difficulties. The pilot survey ran from February 14 to February 28, 2019. The National survey followed, and the data from this National survey is discussed in this paper. The national survey ran from March 5 to June 14, 2019. The survey was conducted online using LimeSurvey (v2.01). Eligibility to participate was determined by the initial questions in the survey.

## Supporting information

Supplementary Table 2: Survey Questions

## Data sharing

Full data sets will be shared upon request and with the approval of the Federation University of Australia Human Research Ethics Committee.

## Acknowledgments

Katherine Christian is supported by an Australian Government Research Training Program (RTP) Fee-Offset Scholarship through Federation University Australia. MRD is supported by a NHMRC Fellowship (APP1130013). The Translational Research Institute is supported by Therapeutic Innovation Australia (TIA). The Australian Government supports TIA through the National Collaborative Research Infrastructure Strategy (NCRIS) program. The Authors would like to thank Dr. Kathryn Futrega for critical discussion and figure design.

