## Supplementary Table 1 for "Survey of Australian STEMM Early Career Researchers: job insecurity and questionable research practices are major structural concerns"

**Supplementary Table 1.** Select comments from respondents to Survey Question 73; *How does your job as an early-career researcher meet your original expectations?*

| Quote number | Specific response |
|---|---|
| 1 | Love the work, but out of PhD the funding applications and extra responsibilities were a big learning curve |
| 2 | Grant funding is extremely difficult to obtain and it is very demoralising |
| 3 | I think I expected as a senior postdoc, greater support from supervisor on the research front e.g. inclusion in grants and publications that I do not lead. In contrast, it has been a case of supervisor offering opportunities in teaching and industry-research, which I have embraced and do well, but retrospectively supervisor blames me for wasting research time when it was it fact on his suggestion that I take those opportunities to diversify and develop a range of skills in the first instance! |
| 4 | My actual day-to-day work far exceeds expectations; I thoroughly enjoy the challenge and freedom of research inquiry, and feel lucky to be able to do this type of work. However, job insecurity is a far greater problem than I had ever anticipated. Though only 3 years post-PhD I have a strong track-record of research output and competitive funding successes. Despite this, there are no foreseeable opportunities to gain a more secure position. I wish to remain at the same institution, or at least within this state because of family and my partner's work, and I guess that this has limited my options significantly. The pressure of Fellowship / project funding success is all the more greater as it is the basis of employment for another year. Last year, and particularly with the delay in ARC outcomes announcement, I was physically unwell from the stress. I am petite but have gained approximately 10kgs since finishing my PhD. At one point in my first year post-PhD I had two fixed-term contracts (0.5FTE and 0.2 FTE) and three-four casual contracts. Trying to manage the workload across so many different projects plus teaching was awful. Even so, I do count myself as one of the lucky ones as I am in a better position than most of my peers currently or recently graduating from their doctoral studies. |
| 5 | As stated earlier, I feel the tenure track scheme we have is unethical, exploitative and extremely harmful to staff who try to navigate it. Around that, we have almost no staff - academic or support/professional - and so the people who are left behind following multiple rounds of voluntary redundancies are only able to constantly fail and be overwhelmed.... there is no other choice. Success is not possible at the moment, it is just survival and (waning) hope that things might get better. |
| 6 | so overloaded, colloborations involve ecr doing tons of work but not getting first or last author position, no job security leading to pressure ++ to NOT have work life balance |
| 7 | Only high level research is supported i.e. focused on medical but not paramedical research. Have had no support for my research area either during or after PHD at this workplace. no support or mentoring in funding applications. |
| 8 | I thought that there would be more support and development opportunities. Instead I feel that I have been thrown into the deep end and expected to take on too much. I have also encountered some antisocial behaviour amongst academics, such as senior staff who have attempted to "steal" work I am doing to present as their own. It's cutthroat. |
| 9 | The only postdoc in lab means managing many new staff/visiting fellows/students and training them on top of the admin work, my own research, constant pressure to write manuscripts (my own and any others that I am part of) and being responsible for several inter-department shared equipment. |
| 10 | You know what you're getting into however, the lack of security, support, under appreciation you cannot prepare for. We are all competing for the same funding and limited research real-estate. My supervisor is unethical and a scoundrel who makes this job terrible. She exists to feather her own nest and ECR area commodity to use to this end. I have the support of friends that are researchers, whom without I would have quit. However, I am very good at my research, and this has also saved me. I would still not consider any other job... research is great fun and I am a survivor hahaha ;) |
| 11 | I never thought I would be trapped in this remote, under-supported, under-resourced campus of XXX (redacted). Lacking of equipments and facilities is the main difficulty and no support from higher level to equip this remote campus with equipments makes this even worse. I and my students have to travel 4 hour round trip to the main campus to access some equipments. |
| 12 | I (and most EMCRs that I know) experience challenges managing my workload (which I would like to be lower), but to be honest this is what I always expected in academia! |
| 13 | The pressure to secure large sums of grant funding, publish incredible papers, and supervise students just to stay competitive in the job market is incredibly tough. Especially when it comes down to actually securing a job offer, and not just the interview, it has very little to do with these things anyway. |
| 14 | I was adequately prepared during my PhD about the difficulty of being a post-doc - I wasn't prepared for the amount of admin work that was expected of me along with my normal duties. I expected grant writing etc but just general lab manager roles that are put upon me feel unnecessary and not a good use of my time. |
| 15 | Particularly since returning to work part time after having my 3 children, I spend a lot more time writing grants, ethics, filling out paperwork, general laboratory management, supervising students than doing any actual laboratory based work which I find disappointing as I like the practical work. My funded hours have also decreased with every leave period i've taken due to a lack of funding, so going from 1.0FTE to 0.6 to 0.4 to 0.2 makes doing any practical work difficult. My current 0.2FTE was my decision so I can also study my Bachelor of Midwifery degree. |
| 16 | I did not expect the lack of support from my University, or the amount of internal fighting you have to do to perform research. I expected the constant criticism when submitting grants and papers. I didn't expect the every decreasing funds for research. I did not expect how much more you have to constantly work - always more work, never anything taken away. |
| 17 | Lack of funding is an issue. There is an ever moving goalpost as to what you need to achieve to obtain funding. There is growing expectation to perform "extra curricular" activities related to being a scientist but undertaken in your own time. The hours one needs to work in order to produce results are unsustainable. Work life balance is a joke if you are working in research. The level of remuneration for the work we do is paltry. |
| 18 | I don't know if I could even list them all !- I can't believe I was hired onto a project without funding - and I never even thought to ask if they had it. I have to apply for grants, but no-one has ever shown me how.- I feel utterly unappreciated, unfulfilled and extremely isolated. I used to be part of a large lab group as a PhD. Now I'm stuck largely alone, I won't know if my grants were successful for months - hence there's no feedback on my work, or sense of achievement and I feel very unfulfilled.- I don't know what I will be doing next month, let alone next year or in the next decade. I find the insecurity and lack of structure very uncomfortable. |
| 19 | I did not anticipate the amount of opportunities that become unattainable when trying to manage a young family and a career in science. How does one compete for fellowships again others of similar experience that have not had any career breaks? Also once you return to work, usually part time when still caring for young children, it becomes near impossible to generate first author manuscripts and there is little/no appreciation for middle author manuscripts |

| | |
|---|---|
| 20 | I was constantly told of (and witnessed) the desperate grind to get enough data, gather enough collaborations, and generate a novel enough idea (often outside the researcher's actual interest) in order to get grant funding and justify a position in academia. At the same time, academics portray industry as a place that gets things done like a factory, where you perform a repetitive task like a cog in a machine. It makes all research career prospects coming out of a PhD sound horrible. Instead, I love my job. We get precise goals for projects that need to meet a target to continue, and if they fail a hard decision is made to put time and resources to a new target/project. The research is not allowed to meander and get lost in the weeds. It varies a lot, and I learn a lot. While the detriment is that I'm not always working with the tissue/disease/target that I'd like to, it always keeps the problem-solving fresh. The team is equally positive about what we have, and makes for a great collegiate atmosphere. |
| 21 | I would be willing to make the sacrifices required for this job, in exchange for security. Which is what Tenure Track jobs used to offer. But now there are none, just these temporary teaching positions -- or -- permanent jobs - which inevitably go to people who have several crappy short-term jobs under their belts. This industry is starting to ask too much. |
| 22 | So many of the required skills are not explicitly taught. The path to becoming independent is not clear. Being productive is difficult with multiple competing priorities. Funding is sooooo competitive. |
| 23 | Phd did not train me to be a postdoc - there is no formal training of project management, time management. Its more sink or swim - if you have those skills innately you do better. If you dont you cannot manage. There was not enough information before I started a phd on career development and how much my track record and number of publications in Phd would affect my ability to get a job or progress in academia. |
| 24 | The more senior I get, the less of the work I do that I enjoy. I am passionate about research, but that makes up so little of my position and day-to-day work. Instead, I am focus on project administration, team management, and constantly applying for funding for my salary and my research. I feel everyday that the expectations are greater, but with less and less funding to achieve the goals we are set. It's very unmotivating, negatively impacts my quality of life, and I am definitely considering a move in the short to medium term if it does not improve. |
| 25 | Lack of support/information early on about what funding I can apply for |
| 26 | In an ideal world science and research should be objective but one soon realizes that also in this area relations are everything and the individual performance does not count as much as |
| 27 | The nature of job meets my expectations but the insecurity of job and lack of funding are horrendous that in my wildest nightmare I wouldn't be able to imagine. |
| 28 | Much greater stress than anticipated - impacting mental health negatively. |
| 29 | The difficulties getting funding/lack of funding is worse than i expected, and this directly contributes to the ability to produce high quality research |
| 30 | Currently employed as a post doc, but the amount of time I get for research is very little, and I did not expect the administrative burden and the number of HDR students I supervise - I feel like I often act as a junior group leader in terms of lab management but am not rewarded for this, as my work review primarily focuses on research outputs. Difficulties in balancing my own expectations with my supervisors expectations for me. |
| 31 | Job security is a major issue for me. While I was told it would be hard, I didn't imagine not landing a secure job 7 years after PhD.The institutional work culture is a major concern (bullying, academic misconduct, workplace safety etc., which goes un-noticed) |
| 32 | Harassment and exclusion from authorship on papers after projects get to that point... 18 hour days and working 7 days a week on an ongoing basis and still being told I'm not working hard enough... passing out in the lab and other workspaces in the last 2 months due to health issues resulting from harassment and still being told I'm not working hard enough. Being set up to fail in projects and put in direct conflict with other people's students by my supervisor setting and agreeing to deadlines for my work that could never possibly be met. Being yelled at by my supervisor on a regular basis, being yelled at by his students due to my supervisor lying to the students, being unable to lodge complaints as it's made clear that I will not have my contract continued and will have difficulty finding another job without references if I lodge a complaint. |
| 33 | As a first year post doc I was told I would need to fund my entire salary, despite my KPI being $10k. I have since secured grant funding of approximately $150k, however was told that would not be enough to fund more than a 0.4fte. I was previously working on 0.6fte and would have been satisfied keeping that workload for a period, until I could secure more funding which seemed very likely. The university was not willing to fund that extra 0.2 FTE and 0.4fte is just too low of a FTE. I was already working a 1.0fte on that 0.6FTE pay, which meant I worked 2 days a week for free for 8 months, and working 3 days a week for free did not appeal to me. Additionally for the last 8 months or so at the university, I left 3 weeks ago, I was essentially just employed as a glorified RA. My supervisor became a micro manager who constantly changed priorities, sometimes hourly. I was more micro managed more than I was as a 22 year old graduate. He became very aggressive, swore at me when he thought I hadn't done what he wanted and deliberately undermined my attempts to secure my own funding and work agenda.In addition, research collaborators did not fulfill their publishing commitments, despite placing huge demands on me to ensure every piece of work I did was converted to a publication, while their pubs that I am a co-author on still languish and have not been submitted. As a result I have a few first author pubs, but no co-authored pubs which has limited my employment prospects. |
| 34 | Once I finished my PhD I thought I would have better work life balance. I would finally be able to leave uni for the day and leave my work in the office. This has very much not been the case. I still have to take work home with me. I still feel guilty at home when at home watching tv trying to recover instead of completing some work tasks. Its also harder than doing a PhD sometimes because at least if you needed a day off you just took a day off. Now I'm only someone else's project and there are clear tasks and deadlines that need to be achieved I never feel like I can take a day off to recover. I feel like I always need to be on. I even come into work when im completely run down because I feel I have no other choice. I didn't expect it to be like this. The constant need to be working. |
| 35 | I did not expect that responsibility for running the lab, training students, applying for funding would take so much of the time with no recognition. I also did not expect the isolation |
| 36 | The teaching and supervision part of my job is harder than expected. I would enjoy my job if I didn't have to supervise research students.<br>No time Pressure Students are rude No support Stressful |
| 37 | Balancing research and teaching is much more difficult than I imagined. |
| 38 | I like being a scientist but I didn't realise how mentally exhausting it is to be so worried about job security all the time. It's difficult to make plans for the future when you don't know if you'll be able to get another job after the current one. I also feel that I have to say yes to everything and try to be a jack of all trades to have any chance at the next job in this highly competitive environment. I never feel like I'm doing enough or working hard enough and when I take breaks i feel guilty. |

| | |
|---|---|
| 39 | I have been required to take on a number of small administrative roles, which all add up to a reasonable amount of time every week. |
| 40 | Here in Australia both my wife and I lack visa security. We're on a temporary work visa, and recent changes in skilled migration visa have directly affected us in preventing us from applying for permanent residency due to short-term contracts. For sponsored PR a minimum of 2y contract is required, but the policy of our institution is max. 1y contract. However, without PR it is near-impossible to get a long-term contract, so the dog bites it's own tail... |
| 41 | The job itself meets my expectations, but I have physically fallen apart after working so hard for so long to get here. I have not met my own expectations and simply do not have the physical capacity to work hard anymore. I have developed fibromyalgia from trauma, overwork, poverty and stress over the past 40 years and now that I should be nearing the peak of my career I am crashing. |
| 42 | lack of funding lack of security power imbalance in higher education cognitive bias towards non australian females |
| 43 | Although my job meets my expectations, I find it hard to balance my career and personal life. I thought I could come back home from work and not do any work-related activities at home, but often have to do. Also, I have had to work some weekends due to deadlines approaching. |
| 44 | I have not been able to get sufficient funding to do research. There is little support to undertake research over my teaching role. |
| 45 | Had good opportunities to progress in teaching and leadership but when it came to academic promotion, research was the main criteria even though I was a researcher/teaching scholar. I have great potential in research and have demonstrated this with emerging key publications but unable to progress with research in normal work load. Research time is not protected but research outcomes is what they look for in promotion. A stint of parental leave, time away from teaching and service roles, has helped me to re-evaluate my situation.<br>I expected the increased work load from my postdoc |
| 46 | I didn't expected to be teaching as much as I am now (which is more than expected), and I'm very happy with it. |
| 47 | I love research but my funding is very precarious and there are very few tenured jobs. Those for which I would be suitable (Level B) have much more highly qualified applicants due to paucity of jobs. I do not expect to ever get a tenured job<br>Harder than expected but delighted in how I have risen to the challenges and how it has kept me engaged |
| 48 | The first postdoc was exciting and plenty of opportunities. I didn't meet the expectation of the second postdoc job as I was on parental leaves part of the time. My current (third) postdoc was exciting at first but I find it difficult to be given opportunities to advance my career. Overall, it is a disappointment. Given the current climate, I am lucky to have a job in academia in Australia. |
| 49 | The job itself is great, the fleeting and short-sighted nature of the job is terrible. |
| 50 | Work life balance is extremely difficult. Multiple deadlines, being spread too thin across too many activities (teaching, research, service, admin, etc) makes it very difficult to feel 'on top of things'. Having to catch up on emails/work at home with a young family to care for is extremely stressful. I believe expectations of modern academics and the competitiveness in academia are unsustainable and make it very difficult to juggle personal and professional success/satisfaction. |
| 51 | I loved my PhD and my first postdoc. I had the freedom to produce high impact research without needing to meet criteria for promotion. I am now in my second postdoc without a supervisor. I now have to publish 'X' amount of papers per year and bring in 'Y' amount of funding in order to be considered for promotion. The number of papers I need to publish is such that I have to focus on producing quantity rather than quality, which goes against why I got into science and how I have been trained. I don't like saying this, but I regret going into this line of work. |
| 52 | Lack of leadership and mentoring |
| 53 | Poor job security, casual work, lack of career development, lack of mentoring from my supervisor. |
| 54 | My job is great, it is changing and rewarding. However, I feel like it is getting more and more difficult to maintain a job in research. Peoples expectations are changing as well, we want more out of job security than ever before this is shown through the number of people leaving research for a worse job to maintain job security. |
| 55 | There is more administration that I had expected (paperwork and grant applications) |
| 56 | I have no great complaints about the actual work, I have no problem coming and do the research, I enjoy being a scientist and being in the lab, what I do not like is the lack of supervision, mentoring, guidance, the marked preference I see my boss has for some male members of the lab, the lack of funding opportunities, the lack of job security and that in this job you have to do everything, you have to come up with ideas to apply for grants, to get funding, then you have to do the experiments, you have to do the data analysis, you have to manage the lab, order consumables, manage budget, mentor students without being the official supervisor, then publish the paper but your boss is the main author and when you finish all that, do it again. Being a postdoc is not a job about research, is about being plenty of things more and then be expected to keep on doing more things for the same money and little recognition, without any job security. |
| 57 | Initially, my job met expectations. Over the last few years rising administrational demands and decisions from management have stifled some of my research |
| 58 | There is a lot more administrative work than I anticipated both with teaching and with research. In addition, I struggle stay on top of multiple projects and there doesn't seem to be an easy way to manage this. There doesn't seem to be any software to assist or if there is, no one seems to know about it. I feel like I need a degree in project management. It is also hard to know which opportunities to say yes to and which ones to pass up. Especially when you're desperate for more publications and potential grant applications. |
| 59 | Funding issues and zero job security even if you are very successful, one time you don't get a grant or fellowship then you're out on you're arse |
| 60 | With help from ECR at my Uni I attained industry grant funding, and buying myself out of most of my teaching means that I now have much of the job I want. It does mean doing some research that is not specifically my focus, but it's close and relevant and interesting and meaningful, so I consider myself VERY lucky.<br>I stepped into research career as i thought this would give me the opportunity for lifelong research and innovation. Unfortunately after graduation, the reality is the frustration of an insecure job opportunity. Now i'm just trapped in a strange circle to struggle for a living. |
