## Supplementary Table 2: Survey Questions for "Survey of Australian STEMM Early Career Researchers: job insecurity and questionable research practices are major structural concerns"

**Supplementary Table 1**

**Questions from online survey, and indication of how questions were derived.**

**Eligibility**

1. Do you have a PhD or doctoral qualification?
    - Yes
    - No -Terminate these
    - Currently studying towards this level of qualification –[Terminate these]

2. What is the number of years since completion of your highest degree?
    - 0–1
    - 2–4
    - 5–7
    - 8–10
    - More than 10 years – [terminate these]

3. What is the nature of your employment? (Original question)
    - University, teaching position
    - University, research only position
    - University, combined teaching and research position
    - University and hospital, combined clinical and research position
    - Government research institute (e.g. CSIRO, ANSTO) [terminate these]
    - Research institute
    - Not for profit organisation – [terminate these]
    - Other, please specify

**Demographics**

**4.** What is your gender?
- Male
- Female
- Other
- Prefer not to say

5. What is your age?
- Less than 25
- 25–30
- 31–35
- 36–40
- 41–45
- Over 45

6. Where were you born? If Other please specify your country. (country list derived from the Australian Bureau of Statistics data on HDR student population)
- Australia
- England
- New Zealand
- India
- Italy
- Vietnam
- Philippines
- China
- Nepal
- Malaysia
- Brazil
- Other (please specify)

7. Is English your first language?
- Yes
- No

8. Do you speak a language other than English at home? (If more than one language other than English, provide the one that is spoken most often)
- No, English only
- Yes, Mandarin
- Yes, Italian
- Yes, Arabic
- Yes, Cantonese
- Yes, Greek
- Yes, Vietnamese
- Yes, other (please specify)

9. Where did you receive your PhD or doctoral qualification? (country list derived from the Australian Bureau of Statistics data on HDR student population)
- Australia
- England
- New Zealand
- India
- Italy
- Vietnam
- Philippines
- China
- Nepal
- Malaysia
- Brazil
- Other (please specify)

10. What is your primary research discipline? Select the appropriate Australian FOR code:
    - DIVISION 01 MATHEMATICAL SCIENCES
    - DIVISION 02 PHYSICAL SCIENCES
    - DIVISION 03 CHEMICAL SCIENCES
    - DIVISION 04 EARTH SCIENCES
    - DIVISION 05 ENVIRONMENTAL SCIENCES
    - DIVISION 06 BIOLOGICAL SCIENCES
    - DIVISION 07 AGRICULTURAL AND VETERINARY SCIENCES
    - DIVISION 08 INFORMATION AND COMPUTING SCIENCES
    - DIVISION 09 ENGINEERING
    - DIVISION 10 TECHNOLOGY
    - DIVISION 11 MEDICAL AND HEALTH SCIENCES

**About your family situation**

11. Do you live with a partner or spouse? (Question adapted from [1]; added 2nd and 3rd "yes" answers)
    - Yes –partner of the opposite sex
    - Yes – same sex partner
    - Yes – prefer not to specify
    - No

12. What best describes your partner/spouse's employment status? (Question from [1])
    - My partner works full time in science
    - My partner works part time in science
    - My partner works full time in another sector
    - My partner works part time in another sector
    - My partner is retired or not employed
    - Not applicable

13. Do you have any children under 18 living at home with you? (Question adapted from [1] but added "some of the time")
    - Yes
    - No
    - Some of the time

14. Who is mainly responsible for the care of these children? (Question from [1])
    - I am
    - My partner is
    - We share the care equally
    - Not applicable

15. Are you responsible for the care of any adult due to their ill-health, age or disability? (Original question)
    - No
    - Yes (please explain)

**About your job and work status and workload**

16. What is the name of your institution? (optional)

17. On average, how many hours per week do you work in your workplace, including in field or clinical settings? (Question from [1])
    - o Up to 20
    - o 21-30
    - o 31-40
    - o 41-50
    - o 51-60
    - o 61-70
    - o Greater than 70

18. On average, how many hours per week do you undertake work related to your employment at home? (Question from [1])
    - o Up to 5 hours
    - o 6-10 hours
    - o 11-15 hours
    - o 16-20 hours
    - o 21-30 hours
    - o Greater than 30 hours
    - o Other, please specify

19. What is your employment fraction? (i.e. 0.2 =one day per week) (Question adapted from [1])
    - o 0.2 FTE
    - o 0.4 FTE
    - o 0.5 FTE
    - o 0.6 FTE
    - o 0.8 FTE
    - o 1.0FTE
    - o Other, please explain

20. How would you describe your overall workload? (Question from [2])
    (much too low, about right, too high)

21. In an ideal world, compared to your current workload, how much time would you like to spend on the following tasks? (Question adapted from [2]; with clinical work added)

| | I would like to do more of this | I would like to do about the same | I would like to do less of this | Not applicable |
|---|---|---|---|---|
| Research (active involvement in experiments, data collection, analysis, report writing) | ○ | ○ | ○ | ○ |
| Teaching (including preparation and assessment) | ○ | ○ | ○ | ○ |
| Training and supervision (of students/postdocs/staff) | ○ | ○ | ○ | ○ |
| Clinical work | ○ | ○ | ○ | ○ |
| Fundraising/applying for grants | ○ | ○ | ○ | ○ |
| Administration (paperwork, committees, departmental meetings, etc.) | ○ | ○ | ○ | ○ |
| Service (voluntary services within organization, counselling colleagues/students, etc.) | ○ | ○ | ○ | ○ |

22. Thinking about all of your paid, unpaid, and other research activities since receiving your doctorate/doctorate-equivalent degree, have you: (Select all that apply) (Question adapted from [3]; with co-supervised added)

    - Published papers in conference proceedings?
    - Had articles accepted for publication or already published in a peer-reviewed journal?
    - Submitted articles for publication in a peer-reviewed journal; that were not accepted for publication or published?
    - Published books or book chapters?
    - Been named as an inventor on a patent application(s)?
    - Been awarded peer-reviewed grant funding?
    - Supervised or co-supervised HDR students to completion?

**About Your Job Security and your Funding**

23. In which manner are you employed: (Original question)
    - Full time continuing
    - Part time continuing
    - Full time fixed term contract
    - Part time fixed term contract
    - Contractor / self employed
    - Other (please specify)

24. If you are on a fixed term contract, what is the total length of your [fixed-term] contract? (Question from [1])
    - Less than 1 year
    - 1 to three years
    - More than 3 years (please specify in comment)

A postdoctoral appointment, or "postdoc," is a temporary position awarded in academe, industry, government or a non-profit organization primarily for gaining additional education and training in research. For the next question, please include any position you consider to be a "postdoc" even if your employer did not or does not. Please also count reappointments to the same position as one appointment. (Question adapted from [3]; plus "please specify in Other" added)

25. How many postdoctoral appointments have you had, including your current position if applicable? Select one. If "other" please explain.
    - 1
    - 2
    - 3
    - More than 3 (please specify in Other)
    - Other

26. How is the major component of your salary funded? If Other please explain (Question adapted from [1], plus "Other please specify")
    - I have my own grant
    - I am employed on someone else's grant
    - I am a direct employee
    - I am self employed
    - A combination of two or more of the above
    - Other (Please specify)

27. Does the nature of your research mean you require additional funding in addition to your salary funding to do your research? (Original question)
    - Yes
    - No

28. Please explain how your research costs are funded. If "Other" please explain. (Original question)
    - My salary and research costs are funded together in one grant
    - My salary is funded by one grant/fellowship; my research costs are funded by separate grants
    - I receive funding for my salary only; I do not have separate research costs
    - Other

29. From which of the following did you receive funding in the last three years? (Check all that apply) (Question adapted from [2])
    - Your own institution
    - Government entities in your own country
    - Business or industry (Australian)
    - Private not for profit (Australian)
    - International entities
    - Others (Please specify)

30. Do you currently have adequate funding to allow you to carry out your research? (Original Question)
    - Yes
    - No

**Job satisfaction**

31. To what extent do you agree with the follow statements about your current job? (Question from [1])

| | Strongly disagree | Disagree | Neither agree nor disagree | Agree | Strongly agree |
|---|---|---|---|---|---|
| I am confident my work/contributions are valued by my employer | ○ | ○ | ○ | ○ | ○ |
| I'm confident I can get research grants | ○ | ○ | ○ | ○ | ○ |
| I'm confident I can publish in good journals | ○ | ○ | ○ | ○ | ○ |
| Overall, I find my work rewarding | ○ | ○ | ○ | ○ | ○ |
| I have good career or promotion opportunities | ○ | ○ | ○ | ○ | ○ |
| I have an unreasonable amount of administrative work | ○ | ○ | ○ | ○ | ○ |
| I have good job security | ○ | ○ | ○ | ○ | ○ |
| I have freedom to pursue my own research interests | ○ | ○ | ○ | ○ | ○ |
| I have adequate equipment and resources to do my job | ○ | ○ | ○ | ○ | ○ |
| I am satisfied with my level of income | ○ | ○ | ○ | ○ | ○ |
| I am able to influence decisions that affect me | ○ | ○ | ○ | ○ | ○ |
| I feel safe in my work environment/workplace | ○ | ○ | ○ | ○ | ○ |
| I am satisfied with my workplace's commitment to a diverse and inclusive workplace | ○ | ○ | ○ | ○ | ○ |

32. Thinking about your current workplace to what extent you are satisfied with the following? (Question from [1])

| | Very satisfied | Satisfied | Neither satisfied nor dissatisfied | Dissatisfied | Very dissatisfied |
|---|---|---|---|---|---|
| The criteria for promotion | ○ | ○ | ○ | ○ | ○ |
| The culture of my workplace | ○ | ○ | ○ | ○ | ○ |
| The leadership and management of my workplace | ○ | ○ | ○ | ○ | ○ |
| Opportunities for attending conferences and study leave | ○ | ○ | ○ | ○ | ○ |
| Support for career development/professional development | ○ | ○ | ○ | ○ | ○ |
| Level of resources and equipment to do my job | ○ | ○ | ○ | ○ | ○ |
| Flexibility of working hours | ○ | ○ | ○ | ○ | ○ |

33. If there was one factor you could change that would make a major difference to your levels of job satisfaction what would it be? (Select one ONLY) (Question adapted from [1])
   - ○ Improved working hours
   - ○ More protected time for research
   - ○ Improved leave provisions
   - ○ Improved institutional / organisational culture
   - ○ Improved promotional opportunities
   - ○ Better pay
   - ○ Improved job security
   - ○ Improved mentorship / supervision
   - ○ More family friendly environment
   - ○ Support for career development
   - ○ Other (please specify)
   - ○ None of these. I am very satisfied with my current job

34. Thinking about the last job you left, what was the reason for leaving? (tick all that apply) (Original question)
   - ○ Lack of funding for new contract/further employment
   - ○ Career progression / development
   - ○ The new job is better suited to my interests / skills
   - ○ For better compensation / salary
   - ○ For full-time permanent position
   - ○ Better work-life balance
   - ○ Unhappy with role
   - ○ Looking to relocate / partner was relocated
   - ○ Launch my own business
   - ○ Terminated / made redundant
   - ○ Maternity / paternity leave
   - ○ Retired
   - ○ Personal reasons
   - ○ Unhappy with organisational culture
   - ○ I was subjected to bullying or harassment at work
   - ○ I'd prefer not to say
   - ○ Not applicable
   - ○ Other, please specify

35. To what extent do you agree that your institution both recognises and values the contributions that you make to... (Question from [4])

| | Strongly disagree | Disagree | Neither agree or disagree | Agree | Strongly agree | Don't know | Not applicable |
|---|---|---|---|---|---|---|---|
| Grant / funding applications? | ○ | ○ | ○ | ○ | ○ | ○ | ○ |
| Knowledge transfer / commercialisation activities? | ○ | ○ | ○ | ○ | ○ | ○ | ○ |
| Managing budgets / resources? | ○ | ○ | ○ | ○ | ○ | ○ | ○ |
| Peer reviewing? | ○ | ○ | ○ | ○ | ○ | ○ | ○ |
| Publications? | ○ | ○ | ○ | ○ | ○ | ○ | ○ |
| Public engagement with research? | ○ | ○ | ○ | ○ | ○ | ○ | ○ |
| Supervising / managing research? | ○ | ○ | ○ | ○ | ○ | ○ | ○ |
| Supervising research students? | ○ | ○ | ○ | ○ | ○ | ○ | ○ |
| Teaching and lecturing? | ○ | ○ | ○ | ○ | ○ | ○ | ○ |

**About challenges relating to your work**

36. To what extent have the following characteristics of your workplace culture impacted you or your career advancement? (Question adapted from [2])

VERY SUPPORTIVE- SUPPORTIVE - NEITHER SUPPORTIVE NOR A PROBLEM – NOT SUPPORTIVE/A PROBLEM – VERY UNSUPPORTIVE/ A MAJOR PROBLEM - NOT APPLICABLE

- Level of support from supervisor/manager in applying for promotion
- Guidance received in performance reviews
- Opportunities for professional development
- Opportunities to undertake/complete qualifications
- Access to research funding
- The attitude towards people of my age
- The attitude towards people of my gender
- The attitude towards people of my ethnic background
- The attitude towards people of my sexual orientation
- Availability of informal mentoring

37. To what extent have the following negative characteristics of some workplace cultures impacted you or your career advancement in your workplace? (Original Question)

NEVER A PROBLEM - SOMETIMES A PROBLEM - A SIGNIFICANT PROBLEM

- Inequitable hiring practices
- Harassment based on different power position
- Lack of support from institutional superiors
- Questionable research practices of colleagues within my institution
- Questionable research practices of colleagues outside my institution

38. How many times in your career have you had to change location in order to advance your career? (Question from [1])
    - o I have never changed location
    - o I have moved once
    - o I have moved twice
    - o I have moved more than twice

39. What has been the most significant impact of the move/s? (Question from [1])

40. Have any of the moves involved international relocation? (Question from [1])
    - o Yes
    - o No

41. Please indicate how much you agree or disagree with the following statements about balancing your current professional and personal responsibilities. (Question from [3])

| | Strongly disagree | Disagree | Neither agree nor disagree | Agree | Strongly agree |
|---|---|---|---|---|---|
| You can manage the demands of your position and home life. | ○ | ○ | ○ | ○ | ○ |
| Your work schedule allows you to maintain the overall quality of life you want. | ○ | ○ | ○ | ○ | ○ |
| Your work schedule provides the flexibility to take care of demands at home. | ○ | ○ | ○ | ○ | ○ |
| Your supervisor understands when demands at home interfere with your professional responsibilities. | ○ | ○ | ○ | ○ | ○ |
| Demands at home have slowed down progress on your professional activities. | ○ | ○ | ○ | ○ | ○ |

**About Mentoring and Supervision**

42. A mentor is someone who is there to assist you achieve your personal, academic and career exploration goals. This person is not necessarily your supervisor. Do you have a mentor? (Original Question)
    - Yes
    - No

43. In the last five years have you been mentored in a mentoring scheme in your workplace or through a professional society? (Select as many as apply. If other, please specify). (Question from [1])
    - Yes through a professional society
    - Yes through my institution's formal scheme with a mentor in my current workplace
    - Yes, through my institution's formal scheme with a mentor in another workplace
    - Yes but in an informal arrangement with a mentor in my current workplace
    - Yes but in an informal arrangement in another workplace
    - No
    - Other (Please specify)

44. How beneficial was the mentoring? (Question from [1])
    - Highly beneficial
    - Beneficial
    - Neutral
    - Not beneficial

45. How important to you for career progression are or have been the following types of support from more senior colleagues or mentors? (Question from [2])

| | Very unimportant | Unimportant | Neither important nor unimportant | Important | Very important |
|---|---|---|---|---|---|
| Advice on career decisions | ○ | ○ | ○ | ○ | ○ |
| Introduction to important networks | ○ | ○ | ○ | ○ | ○ |
| Attain a position / job via direct intervention through personal resources (of the supporter) | ○ | ○ | ○ | ○ | ○ |
| Skill training: methodology | ○ | ○ | ○ | ○ | ○ |
| Skill training: fundraising | ○ | ○ | ○ | ○ | ○ |
| Skill training: (scientific) writing | ○ | ○ | ○ | ○ | ○ |
| Skill training: other | ○ | ○ | ○ | ○ | ○ |

46. Over the past two years (or since taking up your current position if that is more recent) have you participated in a formal staff appraisal/performance review? If "Other", please explain. (Question from [4])
    - Yes
    - No
    - Other (please explain)

47. If you have not had a review what is the reason? If "Other", please specify. (Question from [4])
    - You're on probation?
    - You've only recently been appointed?
    - You haven't been invited to do so?

- You haven't arranged this?
- You are not eligible?
- Other (please specify)

48. [If you participated in your institution's staff review/appraisal scheme in the last two years] How would you rate this scheme's usefulness? (Question from [4])

| | Not at all useful | Not very useful | Neither useful or not | Useful | Extremely useful | Not applicable |
|---|---|---|---|---|---|---|
| Overall? | ○ | ○ | ○ | ○ | ○ | ○ |
| For you to highlight issues? | ○ | ○ | ○ | ○ | ○ | ○ |
| In helping you focus on your career aspirations and how these are met by your current role? | ○ | ○ | ○ | ○ | ○ | ○ |
| In identifying your strengths and achievements? | ○ | ○ | ○ | ○ | ○ | ○ |
| In leading to training or other continuing professional development opportunities? | ○ | ○ | ○ | ○ | ○ | ○ |
| In leading to changes in work practices? | ○ | ○ | ○ | ○ | ○ | ○ |
| In reviewing your personal progress? | ○ | ○ | ○ | ○ | ○ | ○ |

49. When you started with your current employer how useful did you find the following? (Question from [4])

| | Not at all useful | Not very useful | Neither useful or not | Useful | Extremely useful | Offered but not taken | Not offered |
|---|---|---|---|---|---|---|---|
| Institutional-wide induction programs | ○ | ○ | ○ | ○ | ○ | ○ | ○ |
| Departmental /Faculty/Unit induction program | ○ | ○ | ○ | ○ | ○ | ○ | ○ |
| The local induction to your current role | ○ | ○ | ○ | ○ | ○ | ○ | ○ |

50. Do you consider yourself a mentor (Question from [2])
    - Yes
    - No

51. If yes, do you have the skills you need to be an effective mentor? (Question from [2])
    - Yes
    - No

**About professional development and training**

52. To what extent do you agree that…(Question from [4])

| | Strongly disagree | Disagree | Neither agree nor disagree | Agree | Strongly agree |
|---|---|---|---|---|---|
| You are encouraged to engage in personal and career development? | ○ | ○ | ○ | ○ | ○ |
| You take ownership of your career development? | ○ | ○ | ○ | ○ | ○ |
| You have a clear career development plan? | ○ | ○ | ○ | ○ | ○ |
| You maintain a formal record of your continuing professional development activities? | ○ | ○ | ○ | ○ | ○ |

53. In which areas have you undertaken, or would you like to undertake, training in these research and academic skills? (Question adapted from [4]; broken into two parts. This and next Question)

| | Undertaken | Not undertaken, I would like to but have no time | Not undertaken, I would like to but not available | This is of no interest to me currently |
|---|---|---|---|---|
| Ethical research conduct | ○ | ○ | ○ | ○ |
| Grant writing | ○ | ○ | ○ | ○ |
| Interdisciplinary research | ○ | ○ | ○ | ○ |
| Intellectual property | ○ | ○ | ○ | ○ |
| Knowledge exchange | ○ | ○ | ○ | ○ |
| Tips for your publishing | ○ | ○ | ○ | ○ |
| Research impact | ○ | ○ | ○ | ○ |
| Research skills and techniques | ○ | ○ | ○ | ○ |
| Teaching or lecturing | ○ | ○ | ○ | ○ |

54. In which areas have you undertaken, or would you like to undertake, training in these generic management skills? (Question from [4])

| | | | | |
|---|---|---|---|---|
| Budget management | ○ | ○ | ○ | ○ |
| Career management | ○ | ○ | ○ | ○ |
| Collaboration and teamworking | ○ | ○ | ○ | ○ |
| Communication and dissemination | ○ | ○ | ○ | ○ |
| Equality and diversity | ○ | ○ | ○ | ○ |
| Personal effectiveness | ○ | ○ | ○ | ○ |
| Project management | ○ | ○ | ○ | ○ |
| Mentoring and being mentored | ○ | ○ | ○ | ○ |
| Public engagement | ○ | ○ | ○ | ○ |
| Supervision of doctoral/masters students | ○ | ○ | ○ | ○ |

**About Career Planning**

55. In general, upon the completion of your highest degree, do you agree you were confident in your career prospects (i.e. obtaining a good job, securing funding, etc.). (5 point Likert scale, strongly disagree to strongly agree) (Question from [2])

56. Whom do you primarily rely on for career development advice? If "Other", please specify. (Select one) (Question from [3])
    - Your current supervisor
    - A previous supervisor
    - A senior colleague in the department or lab from your current position
    - A senior colleague from a previous position
    - Your doctorate/doctorate-equivalent degree advisor
    - No one
    - Other (Please specify)

57. How do you learn about career opportunities that are beyond academia? Select all that apply. (Question from [5])
    - Academic career opportunities are the only ones I am interested in
    - Academia is the only possibility I am aware of
    - My institution provides relevant workshops and resources
    - I cold-contact individuals in jobs that sound interesting
    - My family
    - A professional society that I am a member of provides this information
    - Science publications/jobs boards
    - Journal related to my area of speciality
    - Online resources including blogs
    - LinkedIn, Twitter and other social networks
    - Speaking with people in my lab
    - Speaking with people in my department
    - Scientific conferences
    - Other (please specify)

58. Does your institute have career advisory services for science ECRs? ((Question from [5])
    - Yes, but I haven't had any contact with them
    - Yes, and their offerings have been useful
    - Yes, but their offerings have not been useful
    - No
    - I don't know

59. Which, if any, of the following activities have you done to advance your career? Please select all that apply. ((Question from [5])
    - Attended career seminars and/or workshops
    - Attended networking events
    - Developed my social media profile
    - I have worked out an individualized development plan
    - Discussed my career future with a graduate adviser
    - Discussed my career future with a mentor
    - Discussed my career future with a careers counsellor at my institution
    - Other (please specify)

**About whether you are considering a change in your work**

60. What are your hopes for your research career? Note: Ignore practical constraints! This question addresses what you'd like to do in an ideal world (Question from [6])
    - I'd prefer another job immediately
    - After finishing my current position, I'd look to move away from research
    - I'd like to stay in research for the medium term
    - I'd like to make research my lifetime career
    - Other, (please specify)

61. Within the last five years have you considered any major career or position changes? (Question adapted from [1]; plus "Yes, to move to another area within or outside science Q)
    - No I have not considered any major changes in my job
    - Yes, to take another position in the same field of science within Australia
    - Yes, to take another position in the same field of science overseas
    - Yes, to move to a different position within my field such as management / academia/industry
    - Yes, to move to another area within or outside science. Please specify in comment
    - Yes, to retire

62. Did you take any concrete action to make such changes? If "Other", please specify in the comments box. If you wish to provide further explanation please use the comment box. (Question from [1])
    - No
    - Yes, I applied for another position in the same field in Australia
    - Yes, I applied for another position in the same field overseas
    - Yes, I applied for a different position within my field (e.g. to move to management)
    - Yes, I applied for a position outside my field or outside science. Please specify
    - Yes, I plan to retire within the next five years
    - Other (please specify)

63. Where would you like to be in five years' time? (Original question)
    - In my role and current position
    - In a higher level role, same workplace
    - In a higher level role, different workplace
    - Similar role different workplace
    - Similar role and field overseas
    - In a management role
    - Not working in science; working elsewhere
    - Working in science outside academia
    - Retired, not working
    - Don't know

64. Where do you expect to be in five years' time? (Original question)
    - In my role and current position
    - In a higher level role, same workplace
    - Similar role different workplace
    - Similar role and field overseas
    - In a management role
    - Not working in science; working elsewhere
    - Working in science outside academia
    - Retired, not working
    - Don't know

65. In which area do you expect to work in the long term (say, 10 years +)? (Original question)
    - Career in higher education – primarily research and teaching
    - Career in higher education – primarily research
    - Career in higher education – primarily teaching
    - Career in higher education – primarily research and clinical
    - Other role in higher education

- Research career outside higher education
- Self-employment/running your own business
- Teaching career outside HE
- Self-employed
- Other occupations
- Don't know

66. In which area do you aspire to work in the long term (say, 10 years +)?? Original question)
    - Career in higher education – primarily research and teaching
    - Career in higher education – primarily research
    - Career in higher education – primarily teaching
    - Career in higher education – primarily research and clinical
    - Other role in higher education
    - Research career outside higher education
    - Self-employment/running your own business
    - Teaching career outside HE
    - Self-employed
    - Other occupations
    - Don't know

67. What would be the main reason you would consider leaving a career in research? If "Other" please specify (Original question)
    - Family/carer responsibilities
    - Interpersonal problems with your supervisor
    - Inadequate job security
    - A lack of independent positions available
    - A lack of funding
    - Other (please specify)

**About Career Breaks**

68. Have you have ever taken a period of 6 months or longer away from work anytime during your career? (Question from [1])
    - Yes
    - No
    - Other (please specify)

69. How long was the break that you took? (Question from [1])
    - Up to one year
    - 1 – 2 years
    - 2 – 5 years
    - Greater than 5 years
    - Up to one year, more than once
    - 1 – 2 years, more than once
    - 2 – 5 years, more than once
    - Greater than 5 years, more than once

70. Why did you take time off? (Tick all that apply. If for some other reason, please specify). (Original question)
    - For health reasons
    - To start a family or have more children
    - To care for a sick family member
    - To write papers from your dissertation for publication
    - To travel
    - For additional education
    - You could not find employment
    - For some other reason (Please specify)

71. Which best describes your return to work after the break? (Question from [1])
    - I returned to the same position, full time
    - I returned to the same position and became part time
    - I returned to the same employer but to a different position – full time
    - I returned to the same employer but to a different position – part time
    - I did not return to my position, I returned later to a different employer – full time
    - I did not return to my position, I returned later to a different employer – part time
    - Other (please specify)

72. Do you have a long term health condition or disability that restricts you in your everyday activities and has lasted, or is likely to last, for more than 6 months? (Original question)
    - Yes
    - No

**About the Expectations you had for your Job Satisfaction** (original question)

73. How does your job as an early-career researcher meet your original expectations? If you wish to offer an explanation, please do so in the comment section.
    - my job is much better than I expected
    - my job is better than I expected
    - my job meets my expectations
    - my job has more difficulties than I expected
    - my job has many more difficulties than I expected

74. How do these statements following correspond with your views about the nature of your job? (Question from [7])

    rating from strongly agree to strongly disagree:
    - This is a poor time for any young person to begin an academic career in my field.
    - 'If I had it to do over again, I would not become an academic',
    - 'My job is a source of considerable personal strain'.

75. How would you rate your overall satisfaction with your current job?

(Question from [7])

5 point scale very satisfied to very dissatisfied

76. It is recognised that there are some difficulties for ECRs in working in a research environment in STEMM disciplines. Why do you choose to stay in academia? (Original question)

Open ended, character limitation

**Further Comments**

77. Is there anything you would like to add which has not been covered in this survey? (Original question)

Open ended, character limitation
